# Structural analyses of apolipoprotein A-IV polymorphisms Q360H and T347S elucidate the inhibitory effect against thrombosis

**DOI:** 10.1101/2024.10.21.619523

**Authors:** Aron A. Shoara, Sladjana Slavkovic, Miguel A. D. Neves, Preeti Bhoria, Viktor Prifti, Pingguo Chen, Logan W. Donaldson, Andrew N. Beckett, Philip E. Johnson, Heyu Ni

**Affiliations:** Department of Medicine, University of Toronto, Toronto, ON, Canada; Keenan Research Centre for Biomedical Science, Li Ka Shing Knowledge Institute, St. Michael’s Hospital, Toronto, ON, Canada; Canadian Blood Services Centre for Innovation, Toronto, ON, Canada; Royal Canadian Medical Service, Ottawa, ON, Canada; Department of Physiology, University of Toronto, Toronto, ON, Canada; Toronto Platelet Immunobiology Group, Toronto, ON, Canada; Department of Laboratory Medicine and Pathobiology, University of Toronto, Toronto, ON, Canada; Department of Biology, York University, Toronto, ON, Canada; Department of Chemistry, York University, Toronto, ON, Canada

**Author notes:** **To whom correspondence should be addressed**: Philip E. Johnson, PhD Professor, Department of Chemistry and Centre for Research on Biomolecular Interactions, York University 4700 Keele Street, Toronto, Ontario, M3J 1P3, CANADA.; Heyu Ni, MD, PhD, FCAHS Professor, Department of Laboratory Medicine and Pathobiology, Department of Medicine, Department of Physiology University of Toronto Senior Scientist of Canadian Blood Services Centre for Innovation Room 421, LKSKI - Keenan Research Centre 209 Victoria Street, Toronto, Ontario, M5B 1T8, CANADA. **Statement of prior presentation**. Some of the data in this manuscript has been presented orally at the International Society of Thrombosis and Haemostasis, June 2023.

**Keywords:** Platelet, Apolipoprotein A-IV, Integrin αIIbβ3, Thrombosis, Structural model, Molecular docking, Molecular modeling, Fluorescence anisotropy, Fibrinogen, Receptor structure-function

## Abstract

Apolipoprotein A-IV (apoA-IV) is an abundant lipid-binding protein in blood plasma. We previously reported that apoA-IV, as an endogenous inhibitor, competitively binds platelet αIIbβ3 integrin from its N-terminal residues, reducing the potential risk of thrombosis. This study aims to investigate how the apoA-IV^Q360H^ and apoA-IV^T347S^ mutations affect the structure and function of apoA-IV. These mutations are linked to increased risk of cardiovascular diseases due to multiple single-nucleotide polymorphisms in the C-terminal region of apoA-IV. We postulate the structural hindrance caused by the C-terminal motifs may impede the binding of apoA-IV to platelets at its N-terminal binding site. However, the mechanistic impact of Q360H and T347S polymorphisms on this intermolecular interaction and their potential contribution to the development of cardiovascular disease have not been adequately investigated. To address this, recombinant forms of human apoA-IV^WT^, apoA-IV^Q360H^, apoA-IV^T347S^ variants were produced, and the structural stability, dimerization, and molecular dynamics of the C-terminus were examined utilizing biophysical techniques including fluorescence anisotropy, fluorescence spectrophotometry, circular dichroism, and biolayer interferometry methods. Our results showed a deceased fraction of α-helix structure in apoA-IV^Q360H^ and apoA-IV^T347S^ compared to the wildtype, and the inhibitory effect of dimerized apoA-IV on platelet aggregation was reduced in apoA- IV^Q360H^ and apoA-IV^T347S^ variants. Binding kinetics of examined apoA-IV polymorphisms to platelet αIIbβ3 suggest a potential mechanism for increased risk of cardiovascular diseases in individuals with apoA-IV^Q360H^ and apoA-IV^T347S^ polymorphisms.

## Introduction

Apolipoprotein A-IV (apoA-IV) is a lipid emulsifying protein in plasma that can be interchanged with other apolipoproteins (apoA-I, apoB, and apoE) (1,2). Human apoA-IV is primarily synthesized in the intestine and plays important roles in lipid metabolism, reverse cholesterol transport, glucose homeostasis in plasma, and putative attenuation of thrombosis (2-7). ApoA-IV is reported to exhibit multiple 22-mer amphipathic structures of α-helix bundles and as low as 5% glycosylation (8-10). The gene encoding for human apoA-IV (apoA-IV-1 or apoA-IV^WT^) is reported to have seven single-nucleotide polymorphisms (11,12). The second most abundant apoA-IV polymorphism possesses an ACT to TCT variation at codon 347, which has an allele frequency of 0.22-0.25. This variant encodes a serine to threonine substitution (apoA-IV-1a or apoA- IV^T347S^). The third most common apoA-IV polymorphism with an allele frequency of 0.03-0.12 has a CAG to CAT variation at codon 360, encoding a histidine for glutamine replacement (apoA-IV-2 or apoA-IV^Q360H^) (12-14). Several epidemiological studies have documented that individuals carrying apoA-IV^T347S^ or apoA-IV^Q360H^ allele have an increased susceptibility to coronary heart disease, although the mechanisms behind this correlation have not been adequately explored (15-17).

Platelets, small anucleate cells in the blood, hold key functions in thrombosis and hemostasis (18-20). They also actively contribute to inflammation, immune responses, tumor metastasis, and atherosclerosis (21-24). Integrins are a conserved ubiquitous class of heterodimeric cell surface adhesion proteins. The predominant integrin on platelets is GPIIbIIIa (αIIbβ3), which is essential for platelet aggregation and important for platelet adhesion to the injured vessel wall (18,25-27). Platelet activation leads to the formation of a plug by crosslinking adjacent platelets through the interaction of integrin αIIbβ3 and fibrinogen (Fg), which can recruit additional platelets, coagulation factors, and enhance blood coagulation (25-30). Classically, fibrinogen has been identified as the molecule which mediates platelet aggregation, however, our earlier studies demonstrated that occlusive thrombi can still occur without either or both fibrinogen von Willebrand factor (VWF), suggesting the existence of yet unidentified molecules that bind to αIIbβ3 and contribute to platelet aggregation and thrombosis (26,31-33). Prior studies on apoA-IV have shown that protein-protein interactions depend on a highly conserved N-terminal region and lipid binding depends on key prolines (34,35). In a recent study, we investigated the role of apoA-IV^WT^ in platelet activity and thrombosis, two important factors that contribute to heart attacks and strokes (4,7,19). This study revealed that apoA-IV is a novel ligand of αIIbβ3 via its highly conserved N-terminal region (E1-L38), and negatively charged aspartic acid residues (D5, D13) were potential αIIbβ3 binding sites (4). Also, other structural studies of apoA-IV showed the role of conserved proline residues in shaping the structure and the lipid-binding function of human apoA-IV. Our date demonstrated that apoA-IV is a novel ligand of αIIbβ3 via its highly conserved N-terminal region (E1- L38), and the negatively charged aspartic acid residues (D5, D13) were potential αIIbβ3 binding sites, which are critical for endogenous inhibition of thrombosis (4,5). Furthermore, apoA-IV was able to attenuate platelet aggregation and activation *in vitro* and prevented thrombi *in vivo*. Finally, we demonstrated that circadian rhythms in humans were negatively correlated with platelet aggregation (4).

The present study aims to examine the impact of the three most common apoA-IV variants, namely apoA-IV^WT^, apoA-IV^T347S^ and apoA-IV^Q360H^, on the structural and functional characteristics of this plasma protein and explored a hypothesis that an alteration in the structure apoA-IV variants affect binding to platelet integrin αIIbβ3. The work provides evidence for why individuals with the apoA-IV^Q360H^ and apoA- IV^T347S^ polymorphisms are more susceptible to cardiovascular diseases, providing valuable insights for prevention. This work demonstrates the value of performing comprehensive biochemical studies to understand the role of apoA-IV polymorphisms in cardiovascular disease, platelet-related diseases, and inflammation as well as advancing therapies against these diseases.

## Results

### Circular dichroism showed a change in helicity of apoA-IV polymorphisms

Human apoA-IV polymorphisms (WT, T347S, and Q360H) proteins were expressed and purified as we described elsewhere (4). A series of structural studies on these variants were conducted utilizing circular dichroism spectroscopy to determine the α- helical content of apoA-IV variants (Fig. 1A, S1). ApoA-IV^WT^, apoA-IV^T347S^, and apoA-IV^Q360H^ polymorphisms exhibited two minima at 207 nm and 222 nm indicative of α-helical protein structure (Fig. 1A). The average α-helix contents (fractional helicity) of each variant were calculated and identified that apoA-IV^T347S^ had (48 ± 5)% α-helix content, while apoA-IV^Q360H^ and apoA-IV^WT^ respectively contained (63 ± 5)% and (69 ± 2)% α-helix, demonstrating significant reduction in the mean fractional helicity of apoA-IV^T347S^ polymorphism compared to that in apoA-IV^WT^ and apoA- IV^Q360H^ isoforms (Fig. 1B).

**Figure 1.**
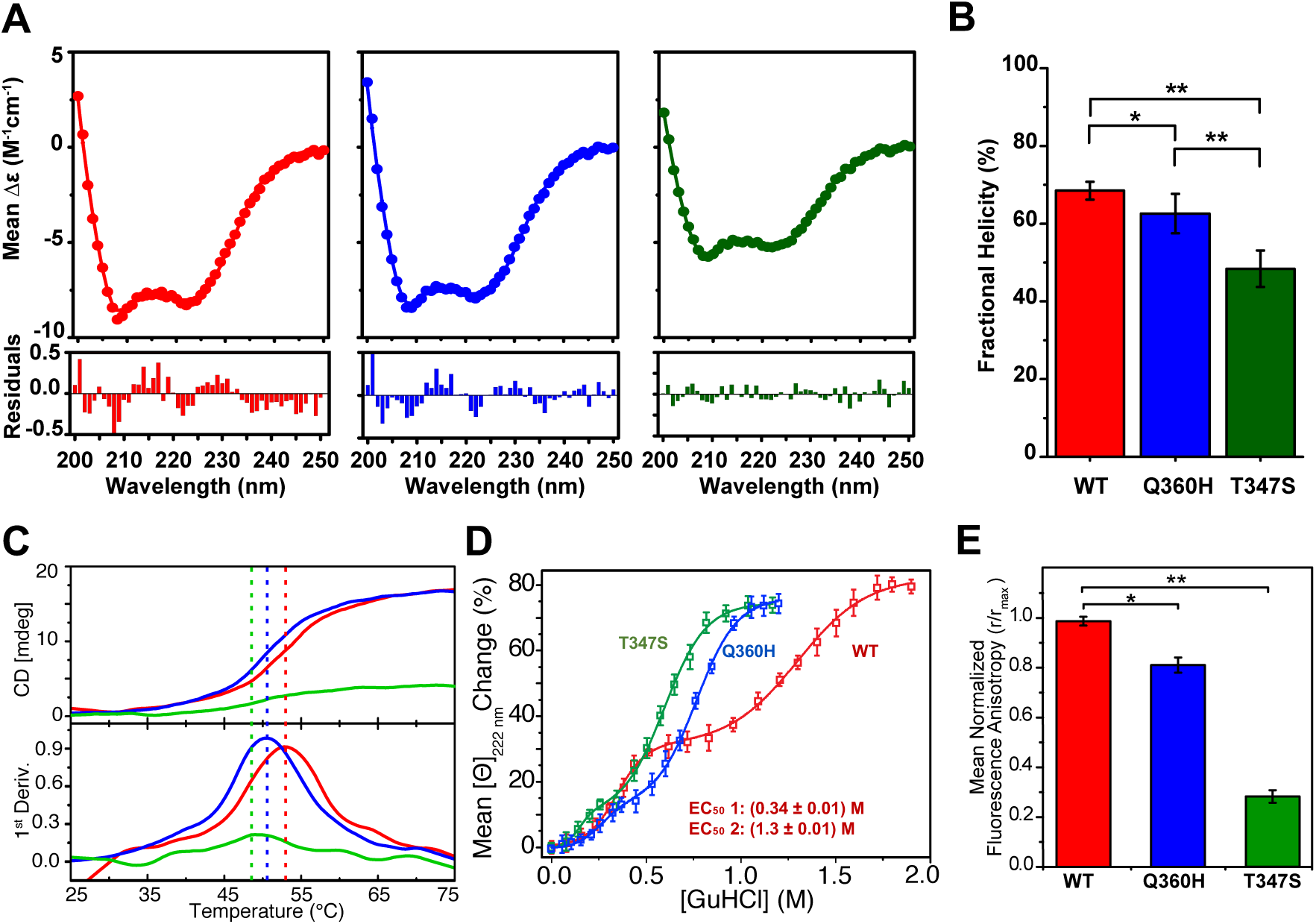
Protein folding and molecular tumbling analyses of apoA-IV polymorphisms. (*A*) shows circular dichroism spectra for the average molar ellipticity versus wavelength of apoA-IV^WT^ (red), apoA-IV^Q360H^ (blue), and apoA-IV^T347S^ (green) in PBS at 37 °C. (*B*) displays a comparison of percentage α-helix content (fractional helicity) of apoA-IV polymorphisms. (*C*) shows the thermal stability analysis of apoA-IV proteins. Doted lines signify T_m_ points of 53.2 °C, 50.8 °C, and 47.6 °C for apoA-IV^WT^ (red), apoA-IV^Q360H^ (blue), and apoA-IV^T347S^ (green) respectively. (*D*) shows protein folding stability analysis of apoA-IV proteins in PBS at 25 °C as a function of GuHCl concentration. (*E*) exhibits molecular tumbling examination using fluorescence anisotropy of FITC-labeled apoA-IV variants in PBS at 37 °C. Fluorescence anisotropy measures the rotational mobility of labeled C-terminus with polarized light. Bar graph demonstrates apoA- IV^Q360H^ tumbles faster than apoA-IV^WT^, and apoA-IV^T347S^ tumbles faster than apoA-IV^Q360H^ and apoA-IV^WT^. Mean normalized fluorescence anisotropy values are subtracted from an unconjugated FITC blank sample. Quantified values are listed in Table 1. * *p* < 0.05, ** *p* < 0.01, N = 20.

**Table 1.** Quantified chemical and thermal denaturation data from CD spectrometry.

| *ApoA-IV* variants | *Chemical stability* |  | *Thermal stability* |
| --- | --- | --- | --- |
|  | *1<sup>st</sup> EC<sub>50</sub>* (M) | *2<sup>nd</sup> EC<sub>50</sub>* (M) | *T<sub>m</sub>* (°C) |
| WT | 0.34 ± 0.01 | 1.3 ± 0.01 | 53.2 ± 0.91 |
| Q360H | 0.26 ± 0.02 | 0.77 ± 0.01 | 50.8 ± 1.1 |
| T347S | 0.15 ± 0.02 | 0.60 ± 0.01 | 47.6 ± 1.0 |

EC<sub>50</sub> values denote GuHCl concentration points at which unfolding occurred. Data presented as mean ± one standard deviation, N = 3.

### Protein folding stability studies revealed denaturation points for apoA-IV isoforms

To test the thermal stability of apoA-IV polymorphisms, we measured mean molar ellipticity, [Θ], at 222 nm in a temperature range of 25-75 °C (Fig. 1C). Using first derivative analyses of the obtained dichroic thermograms, we quantified thermal denaturation points (Tm) for apoA-IV variants as summarized in Table 1. Furthermore, molar ellipticity values of apoA-IV isoforms at 222 nm were measures as a function of GuHCl at 25 °C to examine the chemical stability of apoA-IV isoforms *in vitro* (Fig. 1D). We identified that all three variants showed a biphasic denaturation plot in a dose-response manner with WT demonstrating the most pronounced biphasic denaturation curve. To compare the effect of chemical denaturation, we quantified EC50 values using a biphasic nonlinear regression model (Table 1). The initial transition signifies the disturbance of protein self-association in the solution, while the subsequent transition occurs at a greater concentration of GuHCl, indicating complete protein unfolding. As listed in Table 1, apoA-IV^T347S^ was quantified to have the lowest chemical and thermal denaturation points among the examined variants, and apoA-IV^WT^ showed the highest chemical and thermal denaturation points.

### Fluorescence anisotropy showed molecular dynamics differences in apoA-IV

Molecular dynamics of C-terminal labelled apoA-IV variants were measured *in vitro* utilizing fluorescence anisotropy. Specifically, the molecular tumbling of apoA-IV^T347S^ and apoA-IV^Q360H^ polymorphisms were compared to that of apoA-IV^WT^ in PBS at 37 °C (Fig. 1E). Our results showed statistically significant difference among these three common polymorphisms, with apoA-IV^T347S^ polymorphism demonstrating the lowest average normalized anisotropy value, and apoA-IV^WT^ exhibited the highest average normalized anisotropy value (Fig. 1E). The decreased fluorescence anisotropy observed with apoA-IV^T347S^ and apoA-IV^Q360H^ suggests that these two variants have a faster movement of the C-terminal domain compared to apoA-IV^WT^.

### Dimerized apoA-IV polymorphisms exhibited inhibition of platelet aggregation

The impact of dimerized apoA-IV on platelet function was examined to test whether dimerized apoA-IV protein isoforms (10), which exists in blood, can bridge adjacent platelets. Recombinant apoA-IV protein isoforms were chemically modified and linked using covalent spaces (10), and their effects on ADP-induced human platelet aggregation were investigated. Our findings revealed that the dimerized apoA-IV protein isoforms exhibited a significantly greater inhibition of platelet aggregation compared to the unmodified heterogenic apoA-IV isoforms (Fig. 2A, 2B). Upon analyzing the aggregation findings among dimer apoA-IV variants, we observed that the inhibitory effect was more pronounced in apoA-IV^WT^ dimer in comparison to the apoA-IV^T347S^ and apoA-IV^Q360H^ variants. This trend was also observed with the unmodified heterogenic apoA-IV isoforms (Fig. 2B). This is the first to show that dimerized apoA-IV, which contains two αIIbβ3 binding sites, is still inhibitory (but not bridging adjacent platelets) for platelet aggregation, demonstrating its important physiological role in cardiovascular and other platelet related diseases.

**Figure 2.**
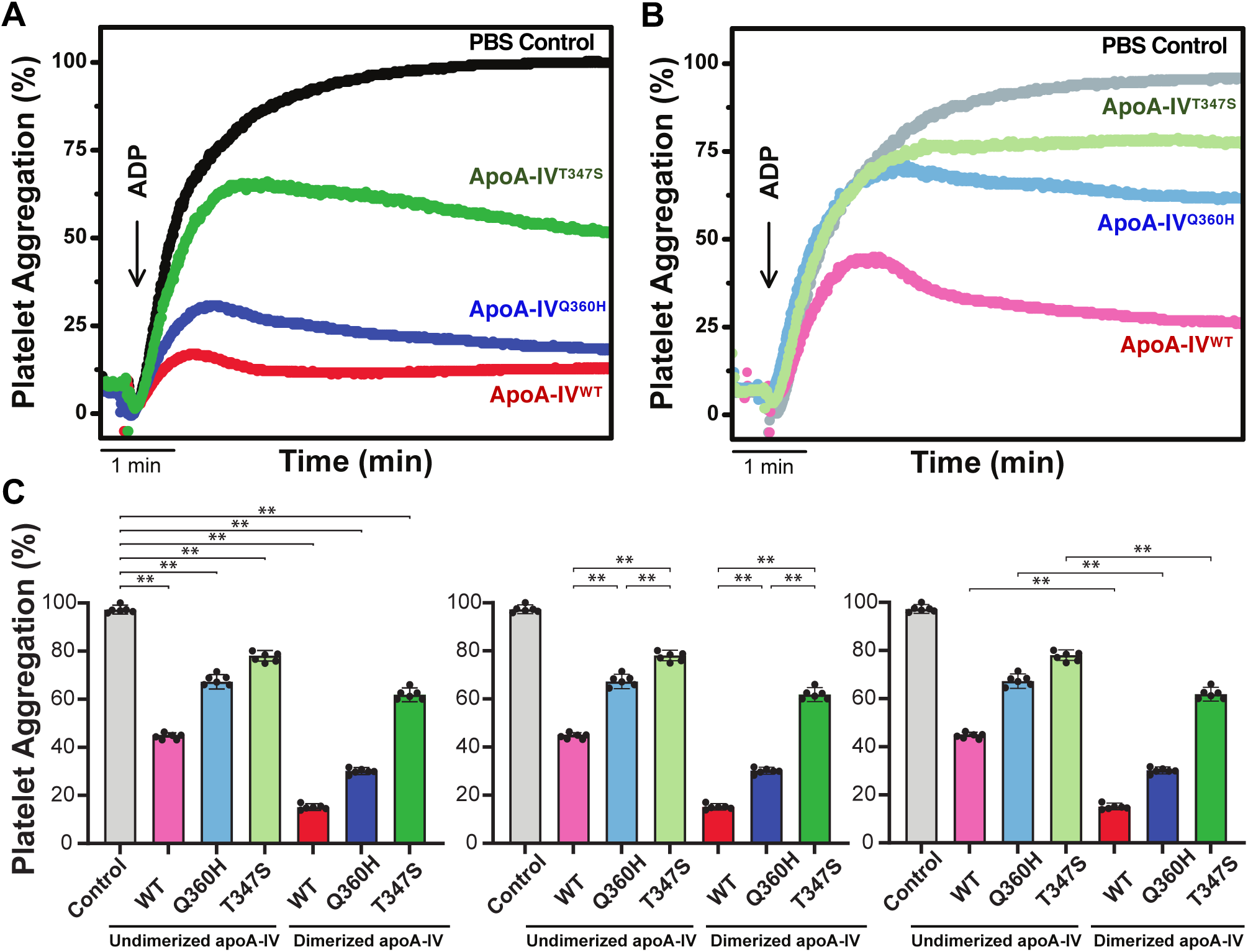
ApoA-IV polymorphisms inhibit human platelet aggregation. ADP-induced human platelet aggregation in PRP with and without the addition of 350 µg/mL (4 µM) apoA-IV^WT^ (red), apoA-IV^Q360H^ (blue), apoA-IV^T347S^ (green), and PBS control (grey) using light transmission aggregometry. (*A*) presents superimposed tracings of platelet aggregation in the presence of dimerized apoA-IV (dark colours). (*B*) shows overlaid tracings of platelet aggregation in the presence of undimerized homogenous apoA-IV (light colours). (*C*) displays bar graphs for comparative statistical analyses of acquired platelet aggregation results. Each bar represents mean ± SD of maximum platelet aggregation. ** *p* < 0.01, N = 6.

### Binding affinity and kinetics studies showed direct binding of apoA-IV to αIIbβ3

To investigate and compare the binding affinities of apoA-IV^WT^, apoA-IV^T347S^, and apoA-IV^Q360H^ polymorphisms with human platelet ligands and receptors, we developed immobilized apoA-IV biosensor probes utilizing an interferometry technique and characterized the kinetics parameters of apoA-IV with activated human platelets at 37 °C using an average weight of (1.6 ± 0.1) ng per activated platelets (36). We found that apoA-IV^WT^ demonstrated the strongest association constant (*ko*n) while apoA- IV^T347S^ variant yielded the weakest *ko*n among examined three variants (Fig. 3A, Table 2). We further tested binding kinetics of apoA-IV variants versus activated αIIbβ3, GPVI, fibrinogen, fibronectin, individually and examined dissociation constant (*K*d) values *in vitro* (Fig. 3B, S2). As presented in Table 2, our results showed that apoA- IV^WT^ exhibited the highest binding affinity with activated αIIbβ3, while the C-terminal mutations (apoA-IV^T347S^ and apoA-IV^Q360H^) displayed weaker binding affinities compared to apoA-IV^WT^ (Fig. 3B). We found no detectable binding signal for apoA-IV interacting with either soluble fibrinogen (Fig. S2A) and fibronectin (Fig. S2B) under the examined *in vitro* conditions. In reverse binding models, immobilized human serum albumin (HSA) and fibrinogen-like 1 protein (FGL1) proteins were tested against (2.0 ± 0.1) µM apoA-IV, and we did not detect binding signals within the experimented conditions. We examined the thermodynamics and dissociation parameters of apoA- IV^WT^, apoA-IV^Q360H^, and apoA-IV^T347S^ at 25 °C utilizing isothermal titration calorimetry (ITC) and quantified dimerization *K*d values as presented in Table 3 (Fig. 4A). The ITC titrations conducted on fibrinogen (Fg) with apoA-IV variants did not yield any quantifiable heat of binding (Fig. S3).

**Figure 3.**
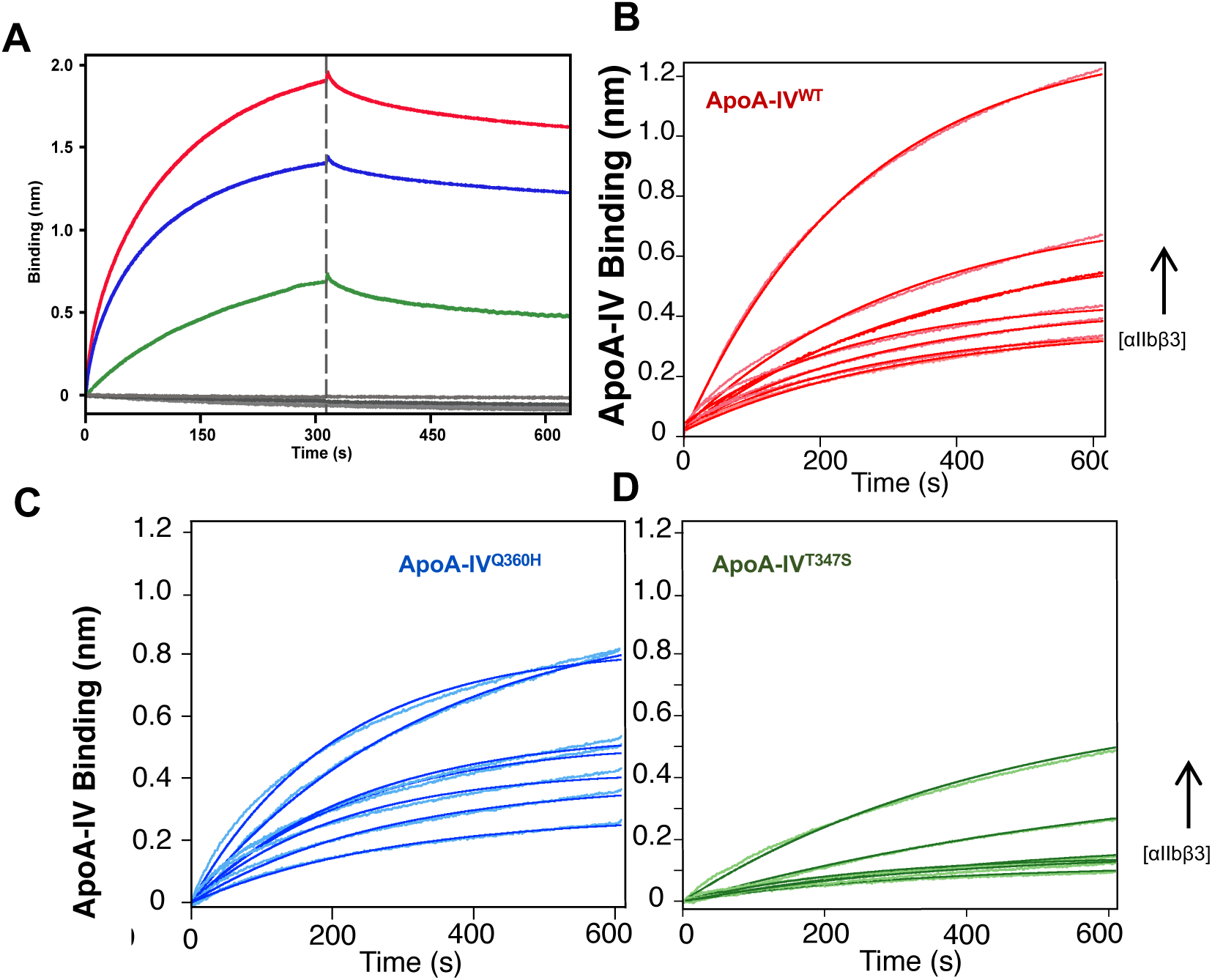
Direct binding affinity quantification of apoA-IV polymorphisms with purified human αIIbβ3. (*A*) displays BLI sensograms for 0.2 µM apoA-IV polymorphisms as a function of activated human purified αIIbβ3 concentration in activation buffer at 1000 rpm, 37 °C (*B*) shows BLI kinetics sensograms for 0.2 µM apoA-IV^WT^, (*C*) apoA-IV^Q360H^, and (*D*) apoA-IV^T347S^ polymorphisms with solutions of gel-filtered platelets in an activation buffer (20 mM Tris, pH 7.4, 137 mM NaCl, 1 mM CaCl_2_, 1 mM MgCl_2_, 1 mM MnCl_2_, 30% (v/v) glycerol) at 1000 rpm, 37 °C (N = 3). Quantified binding affinity values are listed in Table 2.

**Figure 4.**
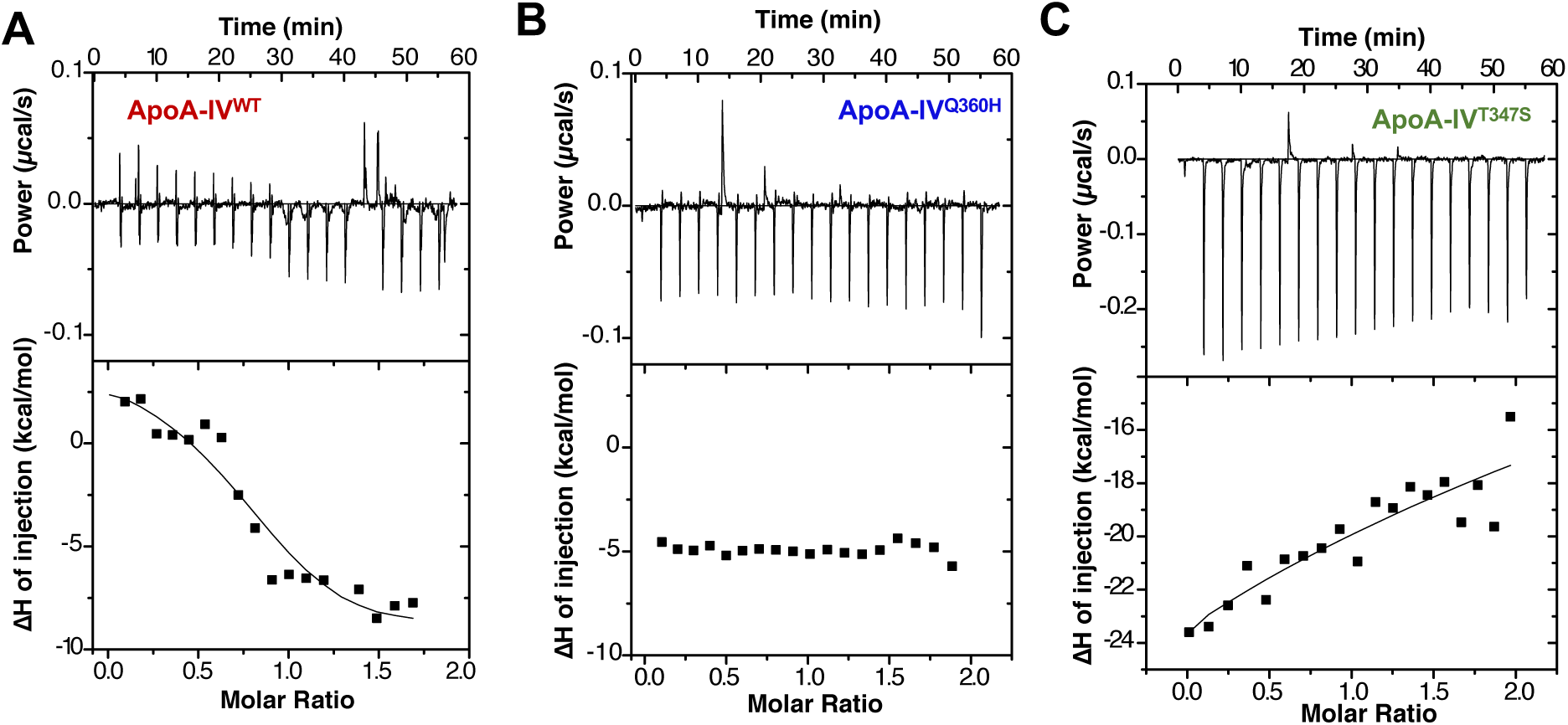
Isothermal titration calorimetry thermograms of apoA-IV polymorphisms. Comparative analysis of multimer dissociation parameters of apoA-IV^WT^ (*A*), apoA- IV^Q360H^ (*B*), and apoA-IV^T347S^ (*C*) in PBS, (N = 3).

**Table 2.**
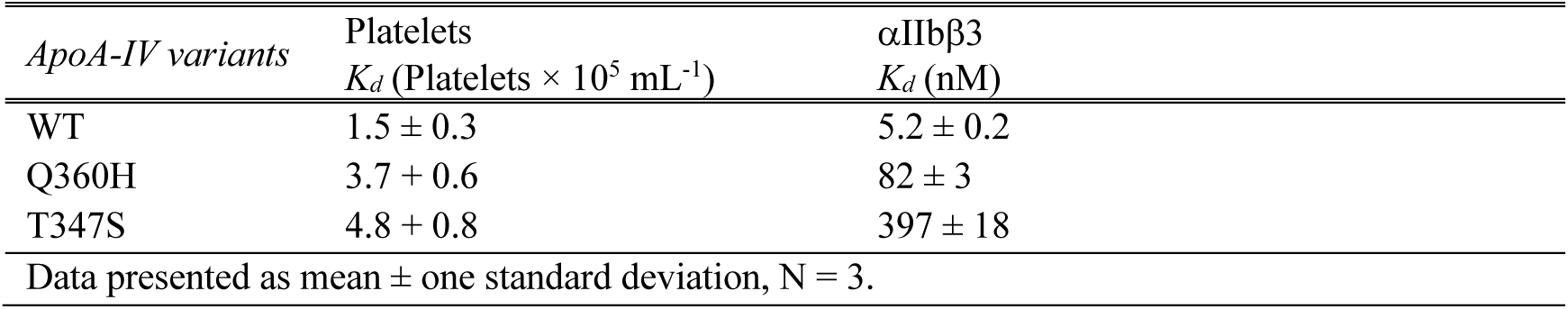
Quantified direct binding affinity data for apoA-IV polymorphisms and human platelets and αIIbβ3 titrations using BLI.

**Table 3.** Dissociation of apoA-IV variants in PBS.

|  | *K<sub>d1</sub>*<br>(nM) | *ΔH<sub>1</sub>*<br>(kcal mol<sup>-1</sup>) | -*TΔS<sub>1</sub>*<br>(kcal mol<sup>-1</sup>) | *K<sub>d2</sub>*<br>(nM) | *ΔH<sub>2</sub>*<br>(kcal mol<sup>-1</sup>) | -*TΔS<sub>2</sub>*<br>(kcal mol<sup>-1</sup>) |
| --- | --- | --- | --- | --- | --- | --- |
| WT | 0.63 ± 0.13 | 2.7 ± 0.6 | -11 ± 1 | 54 ± 17 | -27 ± 3 | 21 ± 4 |
| T347S | 102 ± 51 | -34 ± 1 | -28 ± 1 | - | - | - |
| Q360H | No heat detected |  |  |  |  |  |

Data presented as mean ± one standard deviation, N = 3.

### Light scattering studies demonstrated multimeric conformation of apoA-IV

Dynamic laser light scattering (DLS) and multi-angle light scattering (MALS) techniques were utilized to study the self-association of apoA-IV polymorphisms *in vitro*. Our results showed a heterogenic mixture of apoA-IV multimers and quantified hydrodynamic radii (*R*h) of 2.9 nm, 2.7 nm, and 3.4 nm for apoA-IV^WT^, apoA-IV^Q360H^, and apoA-IV^T347S^ respectively in PBS (Fig. 5A). Our SEC-MALS and non-denaturing PAGE analyses showed significant difference in dimerization properties of apoA-IV variants. We found apoA-IV^Q360H^ eluted a larger amount of monomer conformation than dimer or trimer conformations (Fig. 5B, 5C). We also found both apoA-IV^WT^, and apoA-IV^T347S^ eluted larger amounts of dimer conformation than trimer or monomer. We found that apoA-IV^T347S^ polymorphism eluted the smallest amount of tetramer, or dimer of dimer, conformation among three apoA-IV isoforms (Fig. 5C).

**Figure 5.**
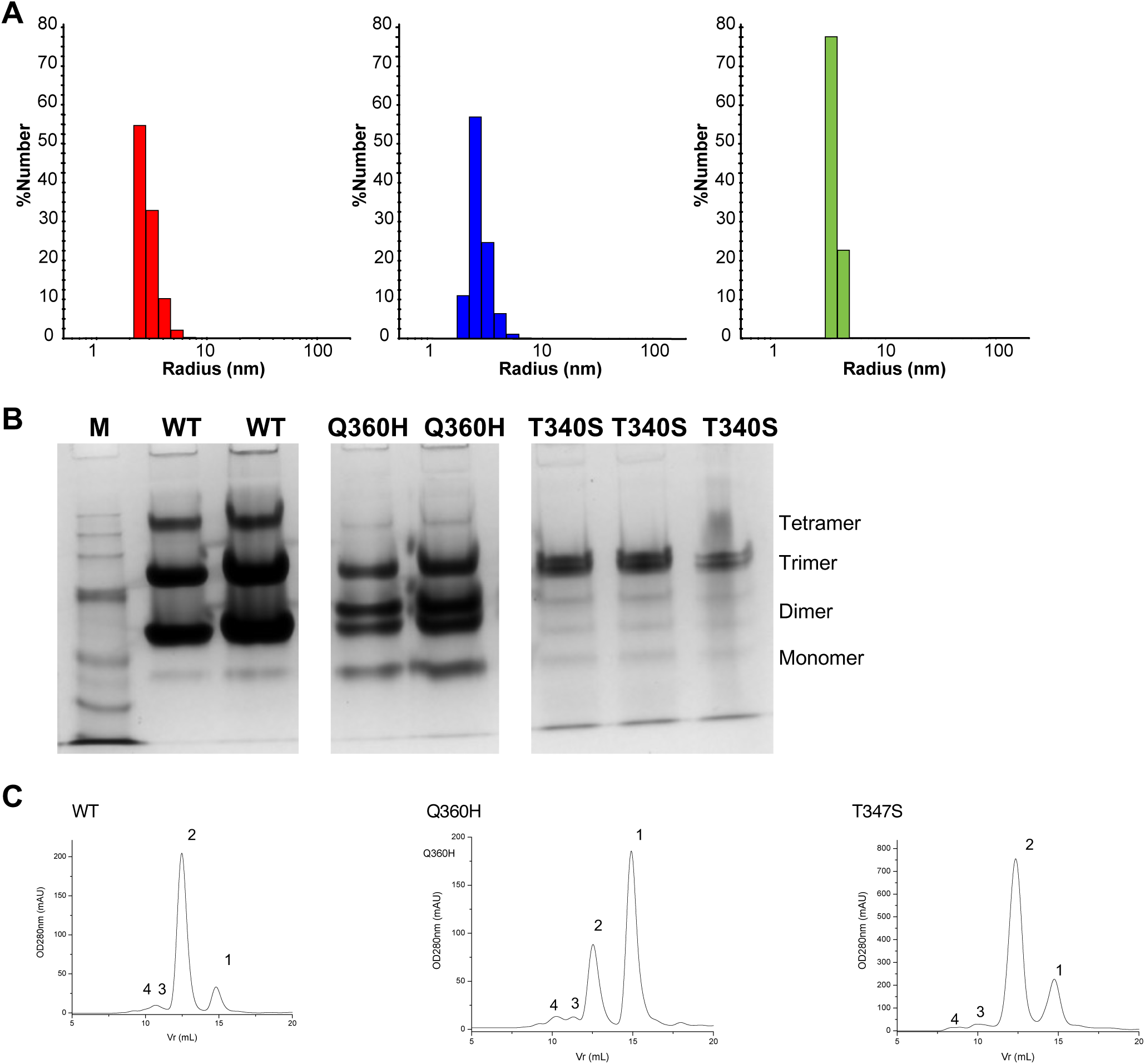
Characterization of molecular size and multimer formation of studied apoA-IV polymorphism. (*A*) shows molecular size analysis of apoA-IV^WT^ (red), apoA-IV^Q360H^ (blue), and apoA-IV^T347S^ (green) using dynamic laser light scattering. (*B*) demonstrates variations in multimer formation properties of apoA-IV on 8% PAGE under non-denaturing pH 7.4 conditions at 25 °C. M denotes protein molecular weight marker. (*C*) shows multiangle light scattering demonstrated multimer diffusion properties of apoA-IV variants in PBS (pH 7.4) at 25 °C. Numeric annotations 1-4 represent eluates corresponding to monomer, dimer, trimer and tetramer, respectively, N = 4.

### Fluorescence quenching studies defined conformational change of apoA-IV

To study the structure unfolding mechanism of apoA-IV polymorphisms through lipid-binding *in vitro*, we measured the intrinsic fluorescence emission of single tryptophan residue (W12) on the N-termini of apoA-IV variants in titrations with DMPC at 37 °C. The intrinsic fluorescence emission of apoA-IV polymorphisms exhibited a dose-dependent rise with the addition of phospholipid DMPC (Fig. 6A). Analyzing the acquired fluorescence intensities and emission maxima, we found that apoA-IV^T347S^, apoA-IV^Q360H^, and apoA-IV^WT^ isoforms yielded a modest shift of (8 ± 3) nm in the emission spectra that was comparable in all three apoA-IV variants, suggesting conformational change in the folded structure of apoA-IV. To further investigate this conformational change mechanism, we studied electron transfer properties of W12 in apoA-IV variants using the intrinsic fluorescence of apoA-IV as a function of two different types of quenchers (sodium iodide and acrylamide) in the absence and presence of DMPC at 37 °C (Fig. 6B, 6C). To prevent the formation of triiodide ion, we added 1 µM sodium thiosulfate in NaI solution and analyzed fluorescence titration isotherms as a function of each quencher concentration and quantified Stern-Volmer constants as summarized in Table 4.

**Figure 6.**
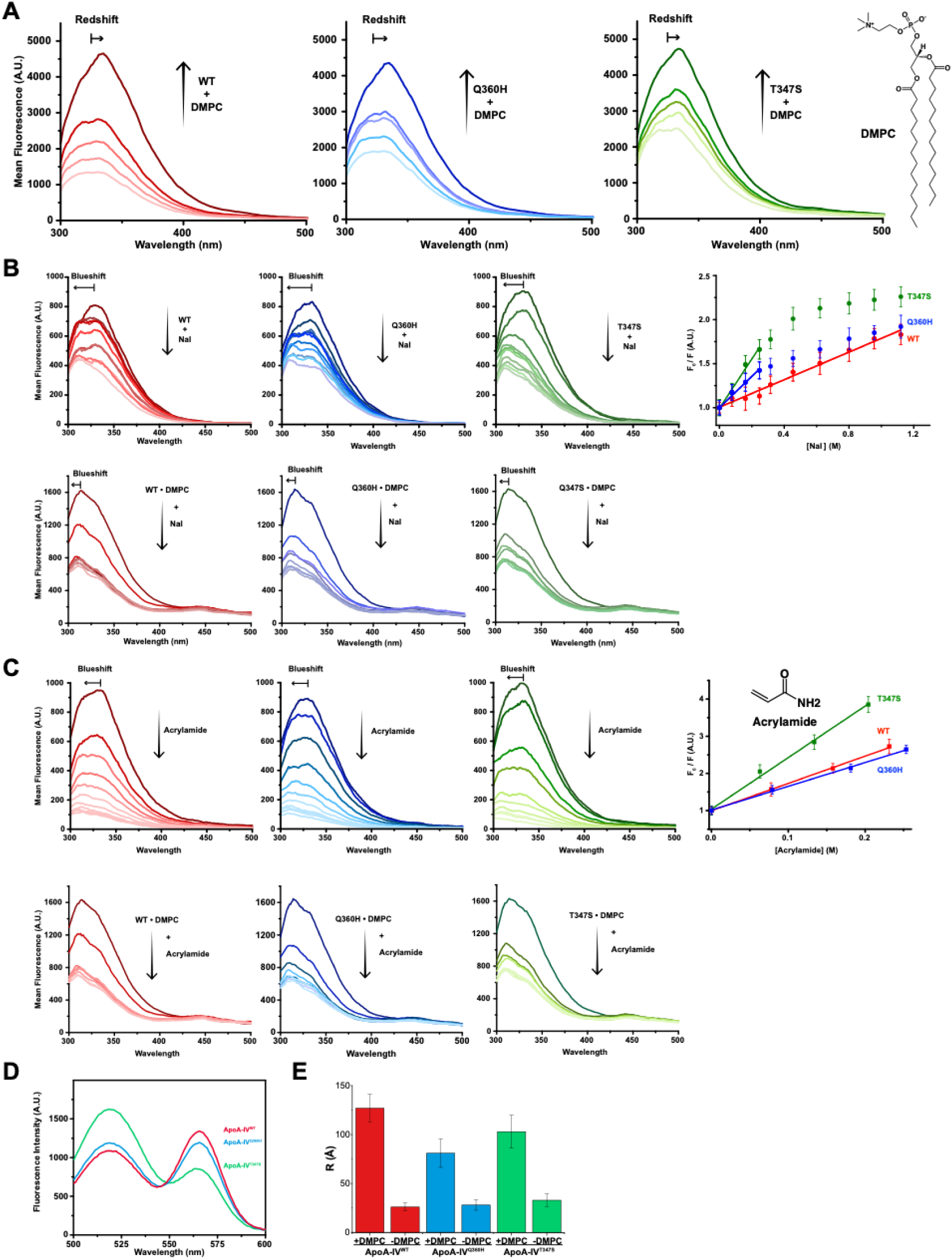
Intrinsic fluorescence enhancement and quenching titrations demonstrating the conformational changes of examined apoA-IV polymorphisms. (*A*) shows fluorescence emission increase of apoA-IV^WT^ (red), apoA-IV^Q360H^ (blue), and apoA-IV^T347S^ (green) as a function of titrations with DMPC. (*B*) exhibits fluorescence emission quenching of apoA-IV polymorphisms bound and unbound to DMPC as a function of sodium iodide concentration. (*C*) shows fluorescence emission quenching of apoA-IV polymorphisms bound and unbound to DMPC as a function of acrylamide concentration. Data acquired in 20 mM HEPES, pH 7.4, 140 mM NaCl at 37 °C. (*D*) Fluorescence emission spectra for FRET analysis of apoA-IV polymorphisms. (*E*) Intermolecular distance comparison of apoA-IV polymorphisms from FRET data without and with 3-fold molar ratio of DMPC.

**Table 4.** Stern-Volmer constants (*K*SV) of apoA-IV polymorphisms quantified from intrinsic fluorescence quenching titrations.

| *ApoA-IV* variants | *Acrylamide*, (M<sup>-1</sup>) |  | *Sodium iodide*, (M<sup>-1</sup>) |  |
| --- | --- | --- | --- | --- |
|  | - DMPC | + DMPC | - DMPC | + DMPC |
| WT | 7.3 ± 0.1 |  | 0.78 ± 0.09 |  |
| Q360H | 6.4 ± 0.2 |  | 1.8 ± 0.1 |  |
| T347S | 13.9 ± 0.7 |  | 2.8 ± 0.3 |  |

Data presented as mean ± one standard deviation, N = 3.

### Fluorescence resonance energy transfer (FRET) reveals spatial structure arrangement of apoA-IV

The spatial structure rearrangement and intramolecular distance of apoA-IV multimers in WT, T347S, and Q360H variants were measured utilizing two FRET techniques *in vitro* at 37 °C. We found that apoA-IV^Q360H^ and apoA-IV^WT^ isoforms emitted higher Förster efficiencies in buffer compared to that of apoA-IV^T347S^ (Fig. 6D), indicating close proximity of the C- and N-termini. We identified that the electron transfer efficiencies were increased immediately after DMPC was added and then decreased and remained steady as apoA-IV variants were mixed with DMPC at 37 °C and incubated for 30 min (Fig. 6D). We measured the intermolecular distance of each apoA-IV variant and found that the distance between C- and N-termini increases from (26 ± 4) Å in a homodimer conformation to (127 ± 14) Å with the addition of phospholipid DMPC, indicating that self-assembly structure of oligomeric apoA-IV was changed and rearranged, where C- and N-termini are farther apart from each other (Fig. 6E).

### Computational structure studies elucidated apoA-IV αIIbβ3 binding interface

*In silico* computational structure modelling was used to analyze the effect of C-terminal mutations of apoA-IV on the structure alignment and binding interface residues with integrin αIIbβ3. While the overall topologies of apoA-IV variants were comparable, we found C-terminal (S336-S376) helical domains of apoA-IV^Q360H^ and apoA-IV^T347S^ exhibited helix-turn conformations whereas apoA-IV^WT^ showed a continuous α-helix structure (Fig. 7A, 7B). Measuring the distance between C-terminal domains from superimposed structures, we calculated apoA-IV^Q360H^ and apoA-IV^T347S^ oriented 20 Å and 26 Å away from apoA-IV^WT^, respectively (Fig. 8A). Analyzing the binding interfaces of apoA-IV with the available structure of the αIIbβ3 headpiece, we computed 41 amino acids of apoA-IV^WT^, including N-terminal D13, interacted with αIIbβ3 (Fig. 8B, 9A) while we found 33 and 27 residues for apoA-IV^Q360H^ and apoA- IV^T347S^, respectively, we did not observe direct contribution of D13 in apoA-IV^Q360H^ and apoA-IV^T347S^ binding αIIbβ3 (Fig. 9A, 9B), indicating apoA-IV variants exhibited distinct binding sites on αIIbβ3 headpiece.

**Figure 7.**
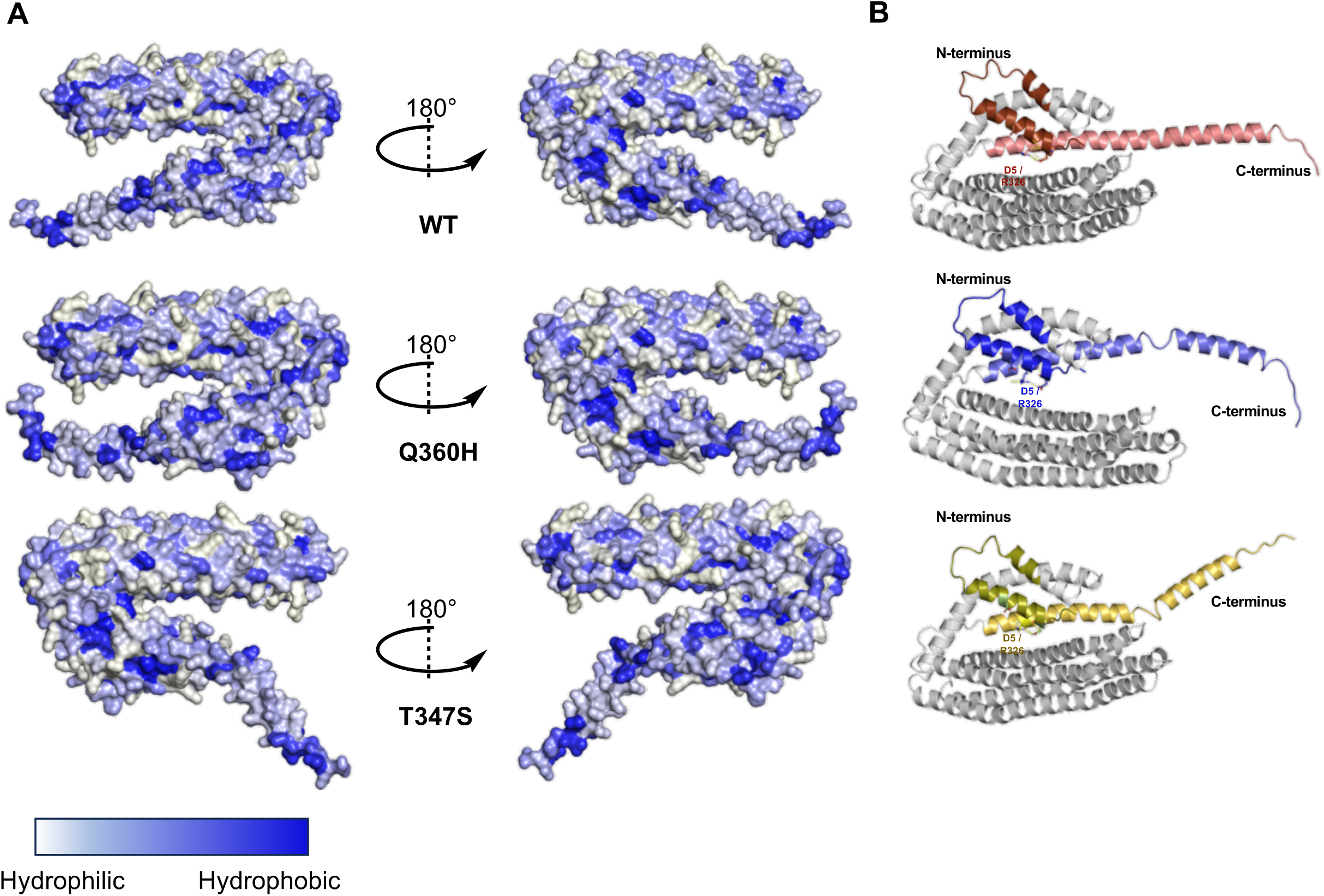
Comparison of overall *in silico* computational structure elucidation of studied apoA-IV polymorphisms in their monomeric conformations. (*A*) demonstrates amphipathic nature of apoA-IV^WT^, apoA-IV^Q360H^, and apoA-IV^T347S^ white colour denotes hydrophilic surfaces, and blue shades signify hydrophilic regions. (*B*) shows differences in spatial orientations of N- and C-termini of apoA-IV^WT^ (red), apoA-IV^Q360H^ (blue), and apoA-IV^T347S^ (olive). Proximity and side chain interaction of aspartic acid (D5) and arginine (R326) residues are denoted on each structure.

**Figure 8.**
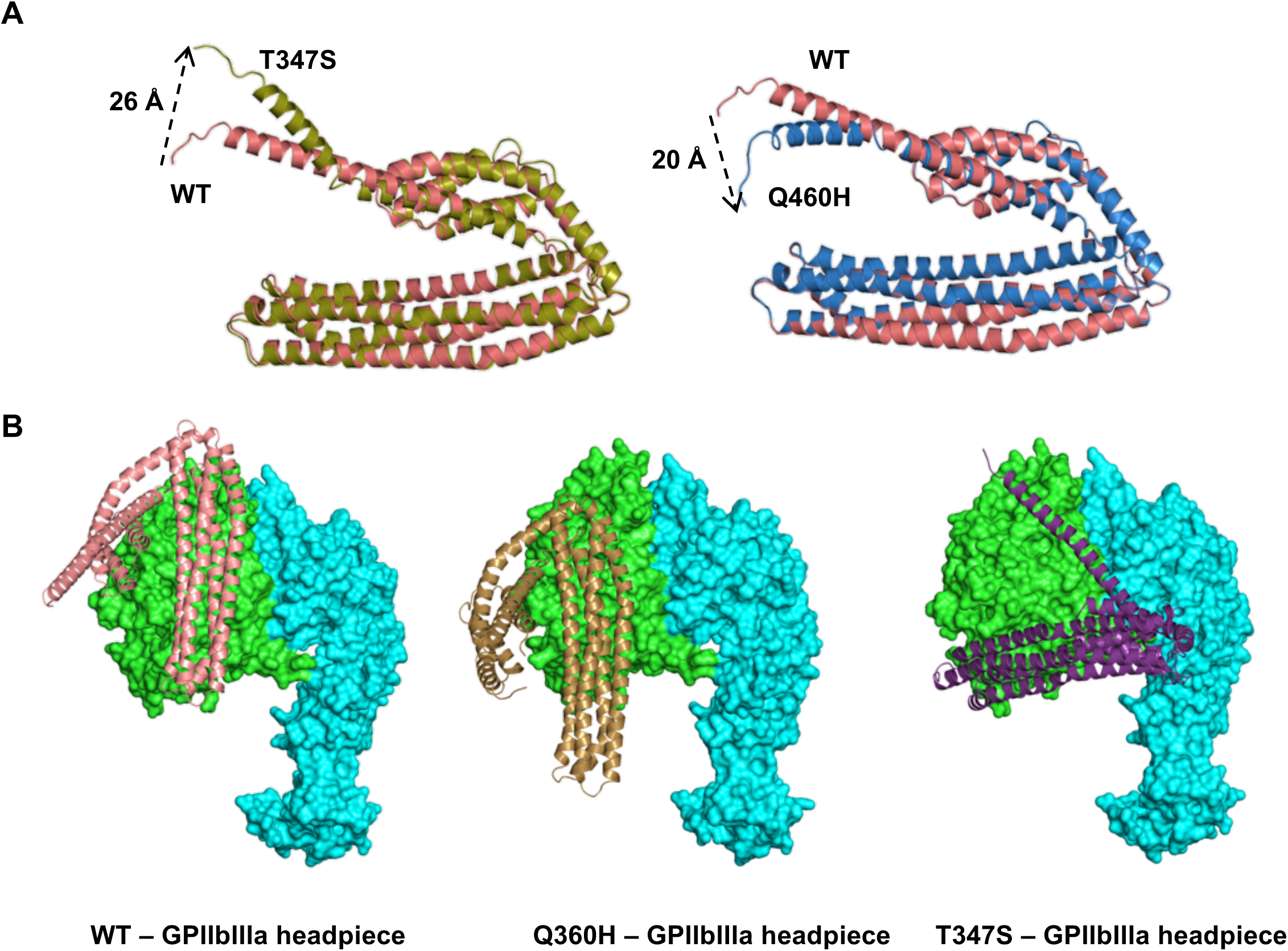
Comparison of C-terminal conformations of examined apoA-IV polymorphisms. (*A*) demonstrates superimposed comparison of apoA-IV^WT^ (red) with apoA-IV^T347S^ (olive) on the left and superimposed comparison of apoA-IV^WT^ (red) with apoA-IV^Q360H^ (blue) on the right. Distance variation in the spatial orientations are denoted on superimposed structures (*B*) shows differences in spatial orientations and interactions of apoA-IV^WT^ (light red), apoA-IV^Q360H^ (gold), and apoA- IV^T347S^ (purple) with the headpiece structure of GPIIbIIIa (αIIbβ3). Protein surface representation of GPIIbIIIa (PTDB: 3ZE2) are shown in green and blue, respectively.

**Figure 9.**
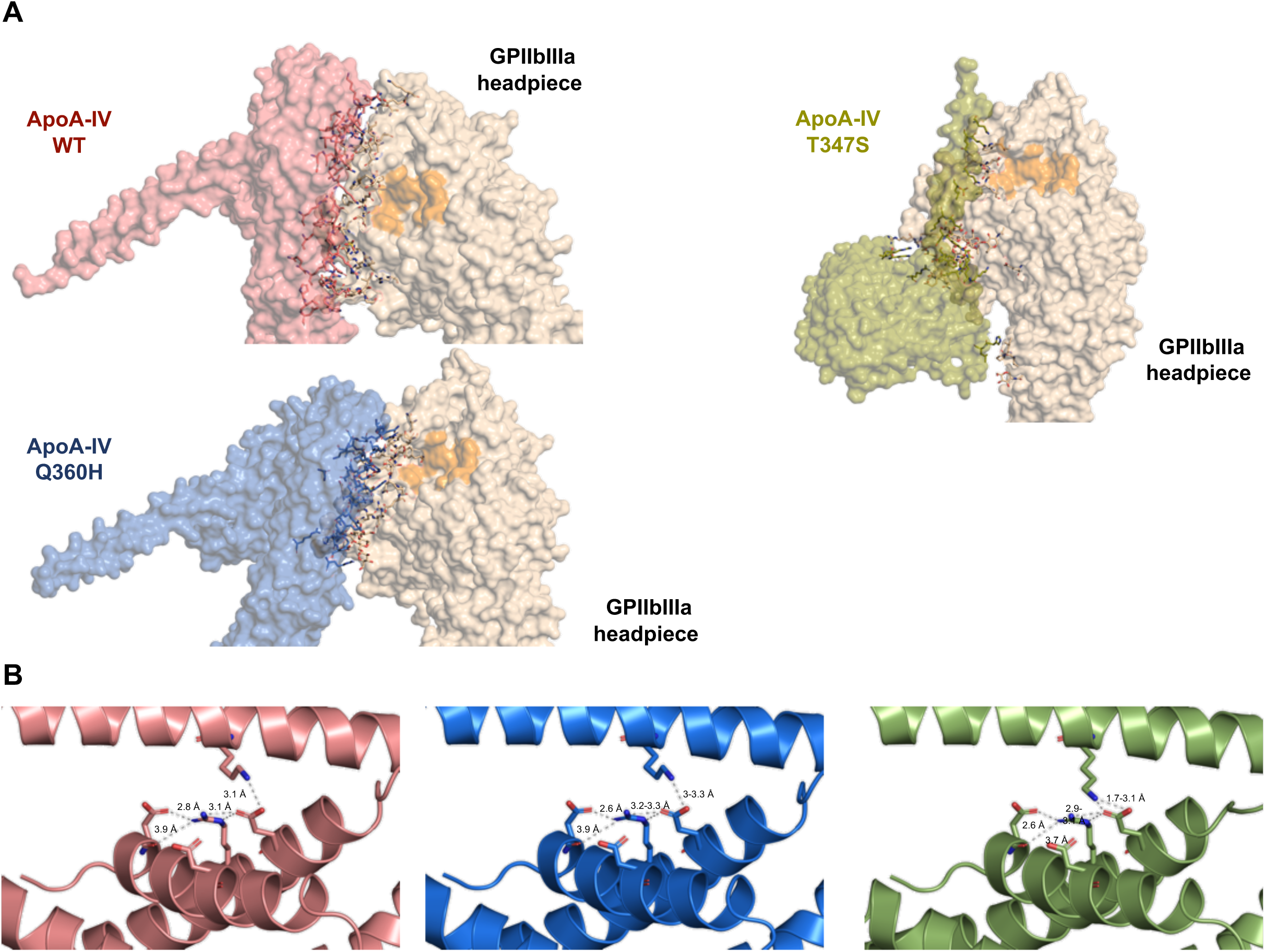
Structural elucidation of direct interactions of studied apoA-IV polymorphisms. (*A*) demonstrates differences in the RGD motif region (orange) on GPIIbIIIa headpiece domain (PTDB: 3ZE2, gold) covered by the binding sites of apoA-IV^WT^ (red), apoA-IV^Q360H^ (blue), and apoA- IV^T347S^ (olive). (*B*) shows zoomed in ribbon representations for intermolecular helix-helix interactions of folded apoA-IV^WT^ (red), apoA-IV^Q360H^ (blue), and apoA-IV^T347S^ (olive) structures.

## Discussion

The impact on the structural stability and functional properties of the three most prevalent apoA-IV protein isoforms were examined utilizing a series of structural and functional assays. Binding kinetics characterizations showed that mutations in the C-terminal region of apoA-IV, specifically apoA-IV^T347S^ and apoA-IV^Q360H^ polymorphisms, can affect the structure of apoA-IV and alter the binding properties of apoA-IV to platelet integrin αIIbβ3. These findings explain our previous observations, wherein N-terminal mutations of D5/D13 abridged the inhibitory effects of apoA-IV^WT^ on platelet aggregation (4,5). In this study, we show that apoA-IV^WT^ and apoA-IV^Q360H^ exhibit higher fractional helicity than that of apoA-IV^T347S^ (Fig 1A). The thermal and chemical stability characterizations demonstrate a significant Tm decrease with apoA- IV^T347S^ and apoA-IV^Q360H^ polymorphisms compared to that of apoA-IV^WT^ (Fig 1, Table 1). These results agree with previous studies, where helix-promoting modifications of apoA-IV favored the higher oligomeric topology and subsequently higher rates of phospholipid binding (35,37,38). The α-helix bundle motif is a key property for lipid-binding protein stability and amphipathic functionality in aqueous environments such as plasma. When apolipoproteins bind to lipids, they form a protective bundle that can be enthalpically favorable to compensate for the unfavorable entropy (39). The apoA-IV^T347S^ variant displays reduced Tm values in contrast with apoA-IV^WT^ and apoA-IV^Q360H^, indicating a less stable protein structure. Thus, apoA- IV^T347S^ is more likely to interact with the charged and hydrophobic phospholipids present in the bloodstream (40).

The fluorescence anisotropy analysis of apoA-IV variants demonstrates significant differences among the three polymorphisms. The molecular tumbling, the rotational movement of molecules around their center of gravity in three spatial axes, and fluorescence polarization and anisotropy values are inversely correlated (41). High fluorescence anisotropy values observed for apoA-IV^WT^ signifies slower C-terminal tumbling to that observed with apoA-IV^Q360H^ and apoA-IV^T347S^ polymorphisms. The order of tumbling observed with the apoA-IV proteins examined agrees with the order of stability yielded from the chemical and thermal denaturation assays (Fig. 1B-1F).

We also studied the impact of apoA-IV on platelet function and tested if dimerized apoA-IV, which physiologically exists in blood, can crosslink platelets. Consistent with our and others earlier studies (4,5,42,43), we show that chemically dimerized apoA- IV^WT^ inhibited platelet aggregation significantly more than dimerized apoA-IV^Q360H^ and dimerized apoA-IV^T347S^. Our results suggest that dimerized apoA-IV polymorphisms do not crosslink platelets and maintain thrombosis attenuating effect as previously described (4,5). The dimerization may be similar to plasma fibronectin (dimerized conformation), in which the two RGD-αIIbβ3 binding sites are close to each other and insufficient to bridge the adjacent platelets, but rather may enhance its inhibitory effect via enhancing the local avidity to block platelet αIIbβ3 integrin (32,44). To the best of our knowledge, this study is the first to present evidence suggesting that dimerized apoA-IV in the blood does not promote, but inhibits, platelet aggregation, and the polymorphisms attenuates this process.

Additionally, immobilized apoA-IV biosensor probes were developed to investigate and characterize the kinetics parameters of apoA-IV variants with activated human platelets. The apoA-IV^WT^ show the strongest binding affinity, while apoA-IV^T347S^ variant had the weakest (Table 2). These results agree with our platelet aggregation results. The binding kinetics of apoA-IV variants with activated αIIbβ3 were tested. The observed binding affinity for apoA-IV^WT^ agrees with our previous published data (4,5). The direct binding affinity of apoA-IV with GPVI, platelet receptor for collagen and fibrin(ogen) interactions (45) were examined as a control and demonstrated no significant binding signals. The absence of binding affinities for apoA-IV variants with soluble fibrinogen, fibronectin, and GPVI signifies the binding specificity and hence the importance of apoA-IV and αIIbβ3 interactions. Our findings agree with previous studies on the main atherogenic constituent of lipid-binding proteins that substantiated the binding properties of lipoprotein(a) with plasma fibrinogen and fibronectin (46,47). It is well-documented that the process of blood coagulation occurs on the surface of phospholipids circulating in plasma (48-50). Existing literature indicates that phospholipids support plasma fibrin polymerization via adsorption of fibrinogen onto phospholipid surfaces (51), thus, we think apoA-IV may regulate fibrinolysis by solubilizing plasma phospholipids. The negatively charged phospholipid bilayers provide a binding platform, where coagulation factors interact and transform proenzyme into active enzymes (28,29,52,53).

We quantified the multimeric diffusion properties of apoA-IV variants and show that apoA-IV^T347S^ variant yields the lowest fraction of tetramer conformation among the apoA-IV variants tested. We attribute this to the increased hydrophobic shielding and negative charge density to the C-terminus of apoA-IV^Q360H^ that can interfere with the domain-swapping dimerization process (10,38,54). A large fraction of monomeric apoA-IV^Q360H^ distribution can also explain weak platelet-binding and small inhibition of platelet aggregation than what we observed with apoA-IV^WT^. We performed the structure alignment of apoA-IV^Q360H^ and apoA-IV^T347S^ variants individually with apoA- IV^WT^ and identified that C-terminal α-helix structure of apoA-IV^Q360H^ had a bent conformation, where it folded closer to the core helical bundle whereas C-terminal α- helix structure of apoA-IV^T347S^ was found to be oriented away from the core helical bundle (Fig. 9). These findings agree with our fluorescence anisotropy analyses, where we detected higher molecular tumbling for apoA-IV^T347S^ and apoA-IV^Q360H^ variants than for apoA-IV^WT^. By comparing the predicted structures of apoA-IV polymorphisms, we were able to identify significant disruptions in the binding interfaces of apoA- IV^T347S^ and apoA-IV^Q360H^ variants with αIIbβ3 compared to that in apoA-IV^WT^-αIIbβ3 binding interface. Our results demonstrate that D5 and D13 residues in apoA-IV have key roles. The D5 residue folds inward and interacts with R326 from the core helix, stabilizing the core bundle and exposing the binding interface. D13 and other 40 residues of apoA-IV^WT^ form a direct interaction platform and shield GRGDSP binding site of fibrinogen and αIIbβ3. ApoA-IV^Q360H^ mutation yields in a binding interface that is reduced to 33 residues, while apoA-IV^T347S^ mutation results in a binding interface that is reduced to 27 residues. Neither apoA-IV^Q360H^ nor apoA-IV^T347S^ binding interface contained D5/D13 residues. The decrease in binding interface residues obtained through our computational analysis is supported by our experimental binding affinity results.

To conclude, we demonstrate significant fractional helicity and structural stability reduction in apoA-IV^T347S^ compared to apoA-IV^WT^ and apoA-IV^Q360H^ isoforms. Our molecular dynamics analysis indicates that the C-terminus of apoA-IV^Q360H^ tumbles slightly faster than apoA-IV^WT^, and apoA-IV^T347S^ tumbles the fastest among examined three variants. The observed trend supports our platelet aggregation, binding affinity, and kinetics results for apoA-IV and αIIbβ3 utilizing our light transmission aggregometry and biolayer interferometry assays. We further elucidate apoA-IV-αIIbβ3 binding interface and demonstrate a biomolecular mechanism for variable functions observed in the most common human apoA-IV polymorphisms: apoA-IV^WT^, apoA-IV^Q360H^, and apoA-IV^T347S^. Our structural analyses provide valuable insights into the structural effects of C-terminal mutations and their potential impact on the function of apoA-IV, which is important for platelets, lipid metabolism, and blood coagulation including fibrinolysis. Notably, as platelets are versatile cells, our data should have broad impact in platelet-related diseases, such as inflammation and immune response, particularly the chronic process of atherosclerosis.

## Experimental Procedures

### Materials

All chemicals and reagents were obtained from Sigma-Aldrich and used as received unless otherwise stated. Buffer solutions were prepared in distilled deionized water with a measured resistivity β18.2 MΘ cm (Milli-Q) at 25 °C and filtered using 0.45 µm filters and sterilized using 0.11-0.22 µm filters. Purified human platelet integrin αIIbβ3 (Innovative Research), purified human plasma fibrinogen, and purified human plasma fibronectin were used as received without any modifications. The protein concentrations were measured using ultraviolet light absorbance at 205 nm, 214 nm, and 280 nm using computed molar extinction coefficient values (Table S1) as described (55). All procedures using human blood samples were approved by the Research Ethics Board of St. Michael’s Hospital, Toronto, Canada.

### Protein expression and purification

Recombinant human apoA-IV^WT^, apoA-IV^Q360H^, and apoA-IV^T347S^ proteins were individually expressed in *Escherichia coli* BL21 (DE3). Full-length human apoA-IV variants were subcloned into the pET30a *E. coli* expression plasmid containing C-terminal 6 × His tag and tobacco etch virus (TEV) protease cleavage. The proteins were expressed by 1 mM isopropyl-β-D-1-thiogalactopyranoside induction (4,8). Harvested cell pellets were lysed using an ultrasonic processor (Sonics) and purified using SuperFlow nickel nitrilotriacetic acid (NiNTA, Invitrogen) columns followed by overnight dialysis in 20 mM Tris buffer, pH 7.4, 137 mM NaCl for binding analyses and separately in PBS (10 mM sodium phosphate buffer, pH 7.4, 137 mM NaCl, 2.7 mM KCl) using (13 ± 1) kDa molecular weight cut-off membranes (Thermo Fisher) at 4 °C. The purification tag was then cleaved using the TEV protease overnight at 4 °C and stopped with 0.1 mM phenylmethylsulfonyl fluoride for 2 hr. The cleaved tag was removed using the NiNTA columns, collecting the eluent purified protein. Purity of proteins were assessed with SDS-PAGE (sodium dodecyl sulfate-polyacrylamide gel electrophoresis) analysis and One-Step Blue (Biotium) staining method.

Electrophoretic mobility shift assays were performed utilizing a non-denaturing PAGE technique at 25 °C as previously described (42,56,57).

### Size exclusion chromatography (SEC)

The size-exclusion chromatography experiments carried out on an ÄKTA Pure Fast Protein Liquid Chromatography (FPLC) system utilizing a Superdex 200 Increase 10/300 column (GE Healthcare). The column was equilibrated with 2 column volumes of PBS running buffer pH 7.4 containing 1 mM ethylenediaminetetraacetic acid (EDTA) prior to each experiment unless otherwise stated. To evaluate oligomeric redistribution of apoA-IV variants, the physiological mid-range concentration (200 µg/mL) of purified recombinant apoA-IV proteins was incubated at 37 °C in running buffer. At different time intervals, 100 μL of each apoA-IV variant was injected onto the column and experiments were run at a constant rate of 0.5 mL/min. The elution process was monitored by measuring the absorbance of eluate at 280 nm. Experiments were recorded and analyzed using the provided Unicorn software package (35,58). The molecular size of the apoA-IV polymorphic multimers were assessed utilizing Multi-Angle Laser Scattering (MALS) with an Infinity-II HPLC (Agilent) and MiniDAWN TREOS and OptiLab T-rEX refractive index detectors (Wyatt). The system was equilibrated in the sample buffer for 18 hours before the first injection and calibrated using 2 mg/mL bovine serum albumin (BSA) standard. Purified recombinant apoA-IV samples (50-100 μL) were injected on to the column and run at a constant rate of 0.5 mL/min. ASTRA software was used to analyze the chromatograms and determine molecular masses (59).

### Chemical modification of proteins

Recombinant human apoA-IV^WT^, apoA-IV^Q360H^, and apoA-IV^T347S^ proteins with a C-terminal cysteine tag were generated in the same manner as described above and conjugated with Alexa Fluor 488 (Thermo Fisher) via a thiol-maleimide reactions. N-terminal apoA-IV samples were conjugated with Alexa Fluor 555 (Thermo Fisher) via N-hydroxysuccinimideester reactions in 20 mM HEPES buffer (N-2- hydroxyethylpiperazine-N-2-ethane sulfonic acid), pH 7.4, 137 mM NaCl at 4 °C as described (60-62). Unconjugated cysteine residues were blocked with 2-fold concentration of N-ethylmaleimide. In separate experiments, 20 µM apoA-IV variants were cross-linked using 0.2 mM bis-sulfosuccinimidyl for 20 h at 4 °C, quenched with 1 M Tris buffer pH 7.4, and buffer-exchanged in 20 mM HEPES, pH 8.1, 140 mM NaCl (10,63). Dimerized apoA-IV proteins were purified using size-exclusion chromatography as described above.

### Platelet preparation and aggregation assays

Human blood samples were drawn from antecubital veins of healthy volunteers after providing informed consent. Human platelet-rich plasma (PRP) and platelet-poor plasma (PPP) were obtained by centrifugation at 250 × g for 8 min and a double spin at 5000 × g for 10 min, respectively, at 25 °C. Gel-filtered platelets were purified from PRP using a Sepharose 2B chromatography column with 5 mM 1,4- piperazinediethanesulfonic acid (PIPES) buffer, pH 7.0, 137 mM NaCl, 4 mM KCl, 5.55 mM glucose as we previously described. After a 5-min incubation, with testing samples or blank PBS (10 mM sodium phosphate, pH 7.4, 137 mM NaCl, 2.7 mM KCl) at 37 ℃. Platelet aggregation was initiated by addition of 5 µM ADP agonist, stirred at 1000 rpm and monitored *in vitro* for a minimum of 8 min using a light transmission aggregometer (Chrono-Log) as we previously described (4,64).

### Biolayer interferometry (BLI) assays

Recombinant His-tagged human apoA-IV^WT^, apoA-IV^Q360H^, and apoA-IV^T347S^ protein isoforms (0.2 µM) were separately immobilized onto hydrated nickel NTA biosensor probes using an Octet RH16 interferometer (Sartorius) and tilted-bottom microplates. The probes were quenched in SuperBlock (Thermo Fisher) and equilibrated in binding buffer (20 mM Tris, pH 7.4, 137 mM NaCl, 30% (v/v) glycerol) containing αIIbβ3 activation divalent salts (1 mM MgCl2, 1 mM CaCl2, 1 mM MnCl2) at 37 °C, 1000 rpm.

The association (*k*on) and dissociation (*k*off) rates were measured to quantify the affinity parameters of apoA-IV variants with αIIbβ3, GPVI-Fc, fibrinogen, fibronectin. The effect of the binding buffer on the threshold of detection was optimized, and raw data were subtracted from the corresponding blank buffer samples. His-tagged recombinant human serum albumin (HSA, Abcam) and His-tagged recombinant fibrinogen-like 1 protein (FGL1, Innovative Research) were used as controls in a reverse binding model. HSA and fibrinogen were immobilized onto separate control probes, and binding signals were monitored with His-tag free apoA-IV^WT^, apoA-IV^Q360H^, and apoA-IV^T347S^ proteins in solution as described above. The acquired data were fit to a global binding model and analyzed to quantify the dissociation constant (*K*d) using the supplied Octet software package as we previously described (65-67).

### Platelet binding kinetics assays

Employing immobilized recombinant apoA-IV^WT^, apoA-IV^Q360H^, and apoA-IV^T347S^ biosensors and the described BLI method, the binding kinetics parameters (*k*on and *k*off) versus activated human gel-filtered platelets were measured in binding buffer (20 mM Tris, pH 7.4, 137 mM NaCl, 30% (v/v) glycerol) containing activation divalent salts (1 mM MgCl2, 1 mM CaCl2, 1 mM MnCl2) at 37 °C, 1000 rpm. Gel-filtered platelets were fixed by mixing platelets in a solution of 2% (v/v) paraformaldehyde (PFA) at 37 °C for 10 min and washed three times between each step (5,66).

### ApoA-IV kinetics assays

The rate of lipid-binding was determined using separate mixtures of apoA-IV^WT^, apoA- IV^Q360H^, apoA-IV^T347S^ proteins, and 1,2-dimyristoyl-*sn*-glycero-3-phosphocholine (DMPC) as a function of time at 37 °C. ApoA-IV protein solutions (0.2 mg/mL) were added to 3-fold concentration of DMPC suspensions in 20 mM HEPES buffer, pH 7.4, 140 mM NaCl. Utilizing 96-well transparent plates (Thermo Fisher) and a Synergy Neo2 (BioTek) microplate reader, the absorbance values at 280 nm were monitored for 30 min (35,68). The acquired data from triplicated experiments were averaged and normalized to the initial absorbance values and analyzed using the first-order decay model as we described (60,66).

### Circular dichroism (CD) spectroscopy

The secondary structures of (2.5 ± 0.5) µM of purified recombinant human apoA-IV^WT^, apoA-IV^Q360H^, and apoA-IV^T347S^ proteins were determined by acquiring CD spectra, immediately after SEC elution, utilizing a Jasco J-1500 spectropolarimeter and 1-mm path length quartz cuvettes in a wavelength range from 195-250 nm at 1 nm/s acquisition rate. The temperature was kept constant at 37 °C using a Peltier controller

(69). The α-helix content (fractional helicity) was calculated from the delta epsilon (M^-1^cm^-1^) values that were deconvoluted from acquired CD spectra in a wavelength range of 200-250 nm using BESTSEL software package (55,70).

### Fluorescence anisotropy assays

The molecular tumbling of apoA-IV^WT^, apoA-IV^Q360H^, and apoA-IV^T347S^ proteins conjugated with C-terminal Alexa Fluor 488 were separately examined by measuring steady-state fluorescence anisotropy (*r*) of Alexa Fluor 488 in PBS at 37 °C. The fluorescence anisotropy of conjugated apoA-IV depends on the ratio of the polarized excitation and emission light, and it is independent of the absolute emission intensity magnitudes (60,71). To ensure the ruggedness of acquired polarization data, the fluorescence anisotropy experiments were repeated utilizing a Synergy Neo2 fluorimeter (BioTek). Using a pair of manual Cary Eclipse light polarizers, the polarized excitation (492 nm) and emission (517 nm) intensities at each titration point were measured and the fluorescence anisotropy (*r*) values were quantified as we previously described (60,67).

### Fluorescence quenching assays

The structural dynamics of apoA-IV variants was investigated exploiting the conformational change of N-terminal single tryptophan (W12) residue of apoA-IV polymorphisms in the presence and absence of DMPC. Tryptophan residues of 10 µM apoA-IV isoforms in 20 mM HEPES, pH 7.4, 140 mM NaCl were excited at 295 nm at 37 °C. The intrinsic fluorescence emission spectra of apoA-IV variants were recorded separately in a wavelength range of 300-500 nm in individual titrations with 8 M acrylamide and 8 Μ ΝaI containing 1 µM Na2S2O3. Equimolar concentrations of N- acetyl-*L*-tryptophanamide (NATA) in the stated experimental conditions were used as control (38,72). The maximum emission intensity change (ΔF) and wavelength shifts (Δλ) were monitored in triplicated experiments. The Stern-Volmer isotherms of the obtained maximum intensities were analyzed as a function of unbound to quenched apoA-IV (F0/F) versus the total quencher concentration. The Stern-Volmer constants (*K*SV) were quantified as previously described (73,74).

### Fluorescence resonance energy transfer (FRET)

Purified recombinant apoA-IV^WT^, apoA-IV^Q360H^, and apoA-IV^T347S^ proteins were separately conjugated with C-terminal Alexa Fluor 488 in one group (donor) and N-terminal Alexa Fluor 555 in another group (acceptor) in 20 mM HEPES buffer (N-2- hydroxyethylpiperazine-N-2-ethane sulfonic acid), pH 7.4, 137 mM NaCl as described above in the chemical modification section applying two FRET techniques: (*i*) C-termini of two apoA-IV batches were labeled with a donor and an acceptor; (*ii*) C- and N-termini of two apoA-IV batches were labeled with a donor and acceptor respectively. Each sample was excited at 488 nm and emission spectra were recorded in a wavelength range of 500-600 nm. The FRET efficiency (*E*) was calculated as *E* = *I*a / (*I*d + *I*a), where *I*a is equal to the integrated fluorescence intensity of acceptor and *I*d is the integrated fluorescence intensity of donor mixed with the acceptor. The intramolecular distance (*R*) between the C-terminus of an apoA-IV molecule and N-terminus of its homodimer was measured as *E* = 1/[1 + (*R*/*R*o)^6^], where *R*o is 70 Å Förster constant for the employed donor-acceptor fluorophores (75,76).

### Thermal and chemical denaturation assays

Utilizing CD spectroscopy, the thermal denaturation assays were monitored from the change in molar ellipticity at 222 nm ([Θ]222) of apoA-IV variants (WT, Q360H, T347S) over the temperature range of 25-75 °C at an increasing rate of 5 °C/min. Dichroic thermograms were subtracted from the corresponding PBS controls, averaged, and normalized. To quantify the thermal denaturation points (Tm), the first derivative of thermograms were plotted as a function of temperature (66,77,78). The thermal denaturation point of unconjugated purified (20 ± 1) µM apoA-IV variants in PBS were examined via intrinsic tryptophan excitation at 280 nm (79). Emission intensities were detected at (325 ± 0.1) nm as a function of temperature increase at 1 °C/min using 2- mm path length quartz cuvettes. The obtained emission intensities were averaged and normalized. To quantify the thermal denaturation points (Tm), the data were analyzed in a same manner as described above for the dichroic thermograms (77,78). The chemical denaturation assays of apoA-IV variants in PBS were examined from [Θ]222 change as a function of guanidine chloride (GuHCl) in a concentration range from zero to 2 M at 20 °C as previously described (37,38).

### Dynamic light scattering (DLS)

The hydrodynamic radii (*R*h) of the apoA-IV polymorphic isotypes were assessed through dynamic light scattering (DLS) utilizing a DynaPro Plate Reader III (Wyatt) (66,80). Minimum volume of 20 μL from purified recombinant apoA-IV variants, with a concentration of (20 ± 1) µM, was introduced into an opaque 384-well clear bottom plate (Corning) and examined at a constant temperature of 25 °C for 5 s/scan. The polydispersity properties of apoA-IV isotypes were determined by analyzing the accumulation of 10-15 trials using the Dynamics software package (66,81,82).

### Isothermal titration calorimetry (ITC)

Thermodynamics and binding parameters of apoA-IV variants were examined utilizing a MicroCal Auto-iTC200 instrument at 25 °C in PBS as we previously described (66,67,83). All experiments comprised of an initial delay of 60 s, first purge injection of 1 μL and subsequent 18 injections of 2 μL titrant, spaced every 180s between every injection. The first point was removed from all data sets due to the difference in injection volume. For dissociation experiments, 43-48 μL of each apoA-IV variant solution were titrated into sample cell containing PBS. Fibrinogen titration experiments with apoA-IV^WT^, apoA-IV^Q360H^, and apoA-IV^T347S^ protein variants were individually performed with 2 µM apoA-IV variant in the sample cell and fibrinogen (29 μM) as titrant. Fibrinogen titration experiments were analyzed with subtraction of the heat of dilution of the titrant. Acquired ITC data were analyzed using Origin 7.0 software provided with the instrument employing the established methods (83,84).

### Computational modeling

Molecular models of full-length human apoA-IV^WT^, apoA-IV^Q360H^, and apoA-IV^T347S^ polymorphisms were computed utilizing a three-track neural network on Colab ProPlus as previously described (66,85,86). The generated secondary structure predictions with high confidence levels were refined and sorted based on the assessed accuracy scores. The structures of apoA-IV variants were compared with available crystal structure of dimeric apoA-IV^WT^ (PTDB: 3S84) (10). The binding interfaces of apoA-IV variants with integrin αIIbβ3 headpiece (PTDB: 3ZE2) (87) were examined utilizing a hybrid algorithm of template-based and *ab initio* free model and analyzed in PyMol (Schrödinger) (88-90).

### Statistical analysis

Data are presented as mean ± standard deviation. Statistical significance was assessed by unpaired two-tailed Student’s *t*-test, one-way analysis of variance followed by Dunn’s test for multiple paired comparisons using GraphPad 10 (Prism). P-value < 0.05 was considered statistically significant (91).

## Data availability

All data generated or analyzed during this study are included in this article (and its supplementary information files).

### Supporting information

This article contains supporting information.

## Acknowledgements

The authors would like to acknowledge the Keenan Research Centre for Biomedical Science Core Facilities at St. Michael’s Hospital (Toronto, ON, Canada). We thank the Hospital for Sick Children (Toronto, ON, Canada) for the use of the Structural & Biophysical Core Facility. We also thank members of the Ni lab, the Johnson lab, and the Donaldson lab for their helpful discussions.

## Author contributions

A.A.S. conceived, developed, devised and performed experiments, analyzed data and wrote the manuscript. S.S. and A.A.S. performed ITC experiments and analyzed data. M.A.D.N., P.B., and P.C. performed initial recombinant protein generation and analyzed data. V.P. and A.A.S. conducted the light transmission aggregometry and analyzed data. L.W.D. and P.E.J. provided lab facility, assisted with structural modeling, and analyzed data. All co-authors assisted in the manuscript writing and editing process. A.N.B., P.E.J., and H.N. are principal investigators, analyzed the data, and edited the manuscript.

## Funding and additional information

H.N. is funded by Heart and Stroke Foundation of Canada (HSFC) GRANT-IN-AID (G-22-0031951); and Canadian Institutes of Health Research (CIHR) Foundation grant (389035). P.E.J. is funded by the National Sciences and Engineering Research Council of Canada (NSERC) Discovery Grant. A.A.S. is a recipient of the Canadian Blood

Services Centre for Innovation Postdoctoral Fellowship and Ontario Graduate Scholarship Awards (York University, Toronto). S.S. and M.A.D.N. are recipients of the Canadian Blood Services Centre for Innovation Postdoctoral Fellowship Awards. V.P. is a recipient of CIHR Graduate Scholarship. The content is solely the responsibility of the authors and does not necessarily represent the official views of the funding agencies.

## Conflict of interest

The authors declare that they have no conflicts of interest with the contents of this article.

### Supporting Information

#### Fibrin clot formation analysis

Freshly prepared human PPP samples were mixed with various concentrations of purified recombinant apoA-IV isoforms or separately with 1,2-dimyristoyl-*sn*-glycero- 3-phosphocholine (DMPC) in 20 mM HEPES, pH 7.4, 140 mM NaCl in 96-well transparent plates (ThermoFisher) and allowed to equilibrate at 37 °C for 10 min. Employing a Synergy H1 (BioTek) microplate reader, the light absorbance at 405 nm was recorded as a function of time. Plasma fibrin coagulation was initiated by dispensing a mixture of 30 mM CaCl2 and 0.1 U/mL thrombin in the stated HEPES buffer conditions. The final volume of each experiment was kept constant at 150 µL. Prior to each scan, the microplate was mixed for 10 s, and the optical density (OD) values were acquired for 30 min (64,92). The acquired data was averaged and analyzed using OriginPro software (OriginLab) and a Boltzmann kinetics model (90,93).

**Table S1.**
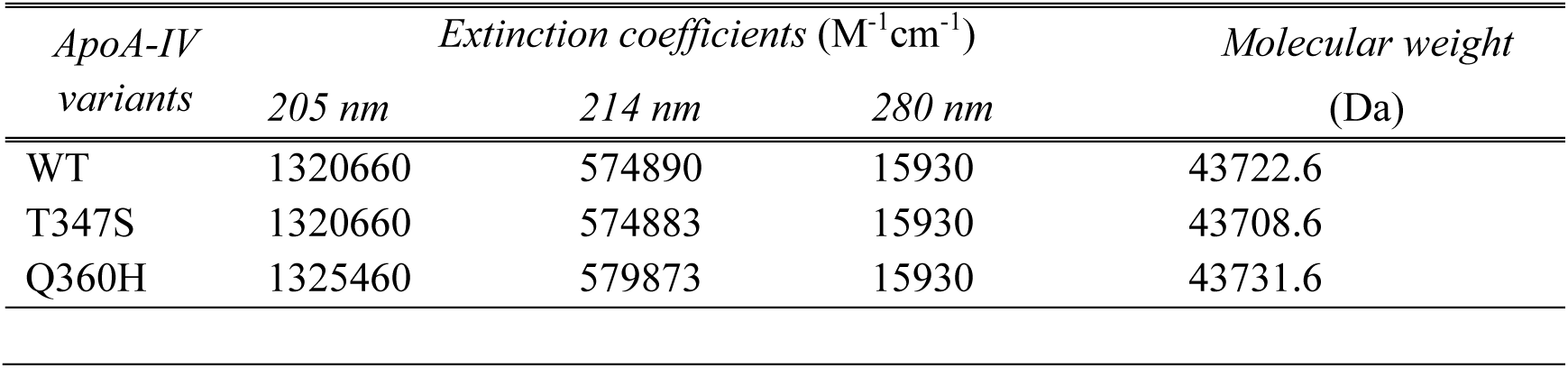
Quantified mass and extinction coefficient values of apoA-IV polymorphisms.

## Notes

### Competing Interest Statement

The authors have declared no competing interest.

