## Supplemental Data for "Structural analyses of apolipoprotein A-IV polymorphisms Q360H and T347S elucidate the inhibitory effect against thrombosis"

***Running title:*** **Apolipoprotein A-IV polymorphisms and thrombosis**

Aron A. Shoara[1-6], Sladjana Slavkovic[2,3,7], Miguel A. D. Neves[2,3,7], Preeti Bhoria[2,6], Viktor Prifti[2,6,7], Pingguo Chen[2,3,6], Logan W. Donaldson[8], Andrew N. Beckett[1-4], Philip E. Johnson[9*], Heyu Ni[1-3,5-7*]

[1]Department of Medicine, University of Toronto, Toronto, ON, Canada; [2]Keenan Research Centre for Biomedical Science, Li Ka Shing Knowledge Institute, St. Michael's Hospital, Toronto, ON, Canada; [3]Canadian Blood Services Centre for Innovation, Toronto, ON, Canada; [4]Royal Canadian Medical Service, Ottawa, ON, Canada; [5]Department of Physiology, University of Toronto, Toronto, ON, Canada; [6]Toronto Platelet Immunobiology Group, Toronto, ON, Canada; [7]Department of Laboratory Medicine and Pathobiology, University of Toronto, Toronto, ON, Canada; [8]Department of Biology, York University, Toronto, ON, Canada; [9]Department of Chemistry, York University, Toronto, ON, Canada.

**Statement of prior presentation:**
Some of the data in this manuscript has been presented orally at the International Society of Thrombosis and Haemostasis, June 2023.

***To whom correspondence should be addressed:**

Philip E. Johnson, PhD
Professor,
Department of Chemistry and Centre for Research on Biomolecular Interactions,
York University
4700 Keele Street, Toronto, Ontario, M3J 1P3, CANADA.

Heyu Ni, MD, PhD, FCAHS
Professor,
Department of Laboratory Medicine and Pathobiology,
Department of Medicine, Department of Physiology
University of Toronto
Senior Scientist of Canadian Blood Services Centre for Innovation
Room 421, LKSKI - Keenan Research Centre
209 Victoria Street, Toronto, Ontario, M5B 1T8, CANADA.

### Figure S1

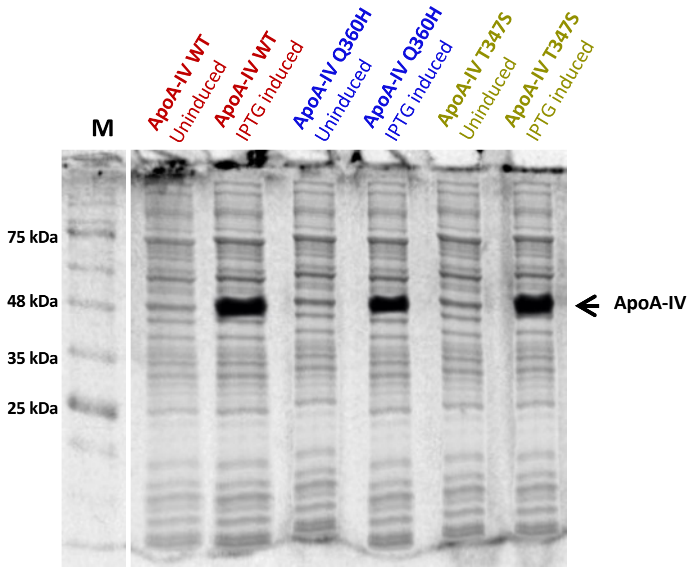


**Fig. S1**. Recombinant expression (*E. coli* BL21-DE3) of apoA-IV protein variants were confirmed using a denaturing and reducing 15% SDS-PAGE. M denotes protein molecular weight marker.

### Figure S2

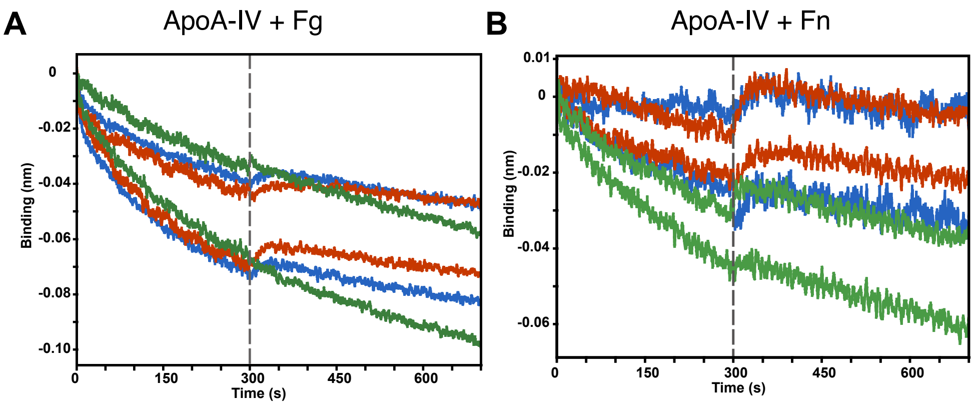


**Fig. S2**. Direct binding affinity quantification of apoA-IV polymorphisms with purified human fibrinogen (Fg) in panel A and fibronectin (Fn) in panel B. Zoomed in BLI kinetics sensograms for 0.2 µM apoA-IV$^{WT}$ (red), apoA-IV$^{Q360H}$ (blue), and apoA-IV$^{T347S}$ (green) polymorphisms in an activation buffer (20 mM Tris, pH 7.4, 137 mM NaCl, 1 mM $CaCl_2$, 1 mM $MgCl_2$, 1 mM $MnCl_2$, 30% (v/v) glycerol) at 1000 rpm, 37 °C (N = 3). No quantifiable binding affinity values were detected.

### Figure S3

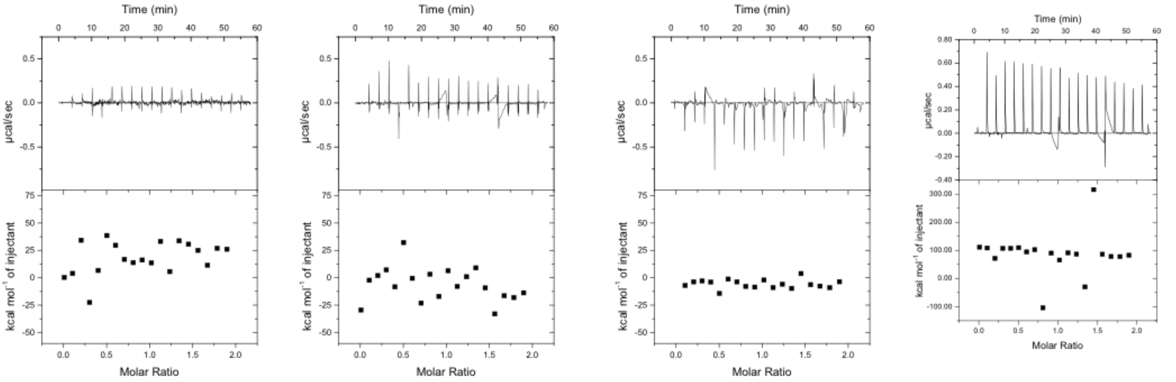


Fg titrating in apoA-IV$^{WT}$ Fg titrating in apoA-IV$^{T340S}$ Fg titrating in apoA-IV$^{Q460H}$ Fg titrating in PBS

**Fig. S3.** Isothermal titration calorimetry thermograms for the direct binding analysis of human fibrinogen (Fg) with apoA-IV$^{WT}$, apoA-IV$^{Q360H}$, and apoA-IV$^{T347S}$ polymorphisms in PBS at 25 °C, (N = 3). Acquired heat signals for the dilution of fibrinogen were subtracted from each titration. No quantifiable binding affinities of Fg and examined apoA-IV protein variants were detected.

### Figure S4

Fibrin formation results

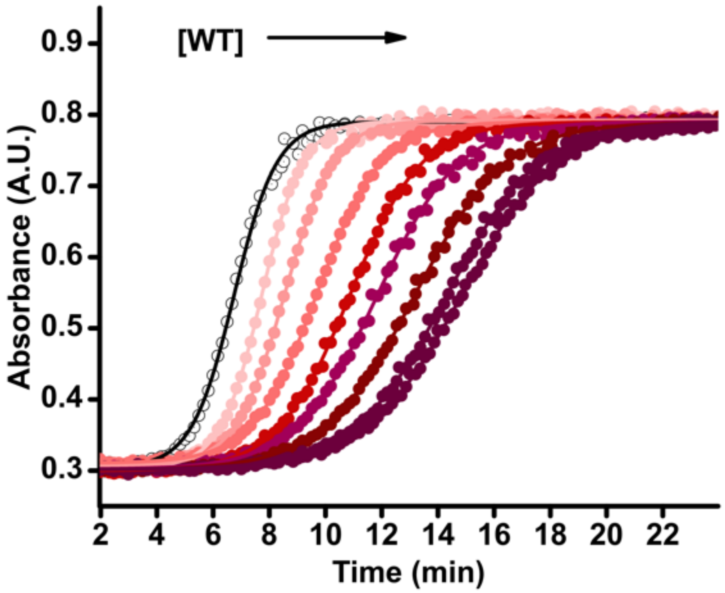


**Fig. S4.** Fibrin network formation analysis of apoA-IV protein variants.
